# The bunyavirus nonstructural protein NSs suppresses plant immunity to facilitate its own transmission by improving vector insect performance

**DOI:** 10.1101/424143

**Authors:** Xiujuan Wu, Shuang Xu, Pingzhi Zhao, Xiangmei Yao, Yanwei Sun, Rongxiang Fang, Jian Ye

**Affiliations:** State Key Laboratory of Plant Genomics, Institute of Microbiology, Chinese Academy of Sciences, Beijing, China; University of the Chinese Academy of Sciences, Beijing, China

## Abstract

Pandemics of vector-borne human and plant pathogens often rely on the behaviors of their arthropod vectors. Arboviruses, including many bunyaviruses, manipulate vector behavior to accelerate their own transmission to vertebrates, birds, insects, and plants. However, the molecular mechanism underlying this manipulation remains elusive. Here, we report that the non-structural protein NSs of *orthotospovirus* (order *Bunyavirales*, family *Tospoviridae*), is a key viral factor that indirectly modifies vector preference and increases vector performance. NSs suppresses the biosynthesis of volatile monoterpenes, which serve as repellents of the vector Western flower thrips (WFT, *Frankliniella occidentalis*) instead of using its known silencing suppressor activity. NSs directly interacts with and relocalizes the jasmonate (JA) signaling master regulator MYC2 and its two close homologs, MYC3 and MYC4, to disable JA-mediated activation of terpene synthase genes. The dysfunction of the MYCs subsequently attenuates host defenses, increases the attraction of thrips, and improves thrips fitness. These findings elucidate the molecular mechanism through which a bunyavirus manipulates vector behaviors and therefore facilitate disease transmission. Our results provide important insights into the molecular mechanisms by which tospoviruses NSs counteracts host immunity for pathogen transmission.

**Author summary:** Most bunyaviruses are transmitted by insect vectors, and some of them can modify the behaviors of their arthropod vectors to increase transmission to mammals, birds, and plants. NSs is a non-structural bunyavirus protein with multiple functions that acts as an avirulence determinant and silencing suppressor. In this study, we identified a new function of NSs as a manipulator of vector behavior, independent of its silencing suppressor activity. NSs manipulates jasmonate-mediated immunity against thrips by directly interacting with several homologs of MYC transcription factors, the core regulators of the jasmonate-signaling pathway. This hijacking by NSs enhances thrips preference and performance. Many human- and animal-infecting members of the *Bunyaviridales* also encode NSs and could manipulate vector behavior to accelerate their own transmission. Therefore, our data support the hypothesis that the NSs protein may play conserved roles among various members of the *Bunyaviridales* in the modification of vector feeding behavior that evolved as a mechanism to enhance virus transmission.

## Introduction

Arthropod-borne viruses (arboviruses) are virulent causal agents of diseases in humans, animals, and plants. Vector behaviors have critical ecological and evolutionary consequences for arboviruses, which rely exclusively on their arthropod vectors for dispersal to (and survival in) new hosts. Therefore, it is of evolutionary significance for an arbovirus to alter its vector’s behavior to facilitate its own transmission. Interestingly, orthotospoviruses, the very few known enveloped plant-infecting viruses, can modify vector feeding behavior to promote their own transmission, as animal-infecting members of *Bunyavirales* [1-3]. Among tospoviruses, *Tomato spotted wilt orthotospovirus* (TSWV) is transmitted mainly by *Frankliniella occidentalis* (Western flower thrips, WFT) in a persistent manner [4-6]. Plants infected with TSWV can affect several vector behaviors, such as biting and host choice behavior [7,8]. However, the underlying molecular mechanism of this conserved manipulation of vector behavior by *Bunyavirales* is still unclear, although host immunity suppression is thought to occur in TSWV-infected *Arabidopsis thaliana* [9].

*Bunyavirales* encompasses nine families of viruses with single-stranded negative-sense RNA genomes. Their genomes are divided into three RNA segments: S, M, and L. In general, the bunyavirus genome encodes four structural and up to two non-structural proteins. The nine bunyavirus families are divided based on their different coding strategies for the additional non-structural proteins, NSm and NSs. The M genome RNA of tospoviruses exhibits an ambisense gene arrangement and encodes an NSm on the viral RNA strand. *Orthotospovirus* NSm protein facilitates the movement of viral ribonucleoproteins from cell-to-cell. NSm of TSWV has recently been identified as the avirulence factor recognized by the product of resistance gene *Sw*-*5b* from tomato (*Solanum lycopersicum* L.) [10]. The S RNA segment is of negative polarity in members of the genera *Orthobunyavirus*, *Orthohantavirus* and *Orthonairovirus*, and ambisense in members of the genera *Phlebovirus* and *Orthotospovirus*. The negative-polarity S RNA encodes the major structural N protein on the complementary RNA strand of the virus, and in certain members of *Orthobunyavirus* and *Orthohantavirus*, the S RNA encodes an additional non-structural protein (NSs) via an overlapping reading frame. The NSs proteins of many bunyaviruses modulate host innate immune responses, and NSs in *Orthotospovirus* functions as a silencing suppressor in both plants and insects [11,12]. These proteins are responsible for establishing systemic infection in plants and for vial transmission by insect vectors [5,13].

Phytohormones such as jasmonate (JA) play vital roles in regulating plant growth and environmental stress responses [14,15]. The core transcription factors in the JA signaling pathway are a class of basic helix-loop-helix (bHLH) proteins, including MYC2, MYC3, and MYC4, whose functions are partially redundant [16,17]. JA-regulated volatile biosynthesis plays an important role in plant-insect vector communication. In addition, the emission of volatiles by many plant species serves as an indirect herbivore defense strategy [18,19]. Several viruses have been shown to modify this process. For instance, begomoviruses inhibit the JA pathway and modify volatile terpene-mediated defense responses against whitefly [20]. A comparison of healthy and TSWV-infected plants showed that thrips feeding normally induces the JA response in plants, whereas viral infection suppresses the JA signaling pathway [21]. However, how TSWV suppresses JA signaling remains elusive, although this virus is thought to hijack the antagonistic relation between JA and salicylic acid signaling [21].

In this study, we identified the NSs protein from thrips-borne TSWV as a viral factor that attracts its insect vector. NSs suppresses the JA signaling pathway in the host plant by directly interacting with MYCs, key regulators of the JA signaling pathway, to reduce host defense responses against thrips; this process functions independently of its virus-silencing suppressor activity. Our results establish a molecular mechanism underlying how TSWV establishes a mutualistic relationship with its thrips vector by targeting the activities of plant MYC proteins.

## Results

### TSWV infection induces a terpene-dependent preference in thrips vector

We first investigated the indirect effect of TSWV infection on the behavioral responses of the vector *Frankliniella occidentalis* (Western flower thrips, WFT). We conducted a two-choice assay between infected and non-infected plants. The model plant *Nicotiana benthamiana*, which is closely related to *S. lycopersicum*, the natural host of TSWV, was used in this study. A group of 50 non-viruliferous WFT were released from the center of the arena between two types of *N. benthamiana* leaves. In this two-choice assay, ~70% of thrips approached TSWV-infected leaves, whereas fewer thrips approached non-infected leaves (Fig 1A), suggesting that TSWV infection indirectly increases the attraction of *N. benthamiana* to thrips vector.

**Fig 1.**
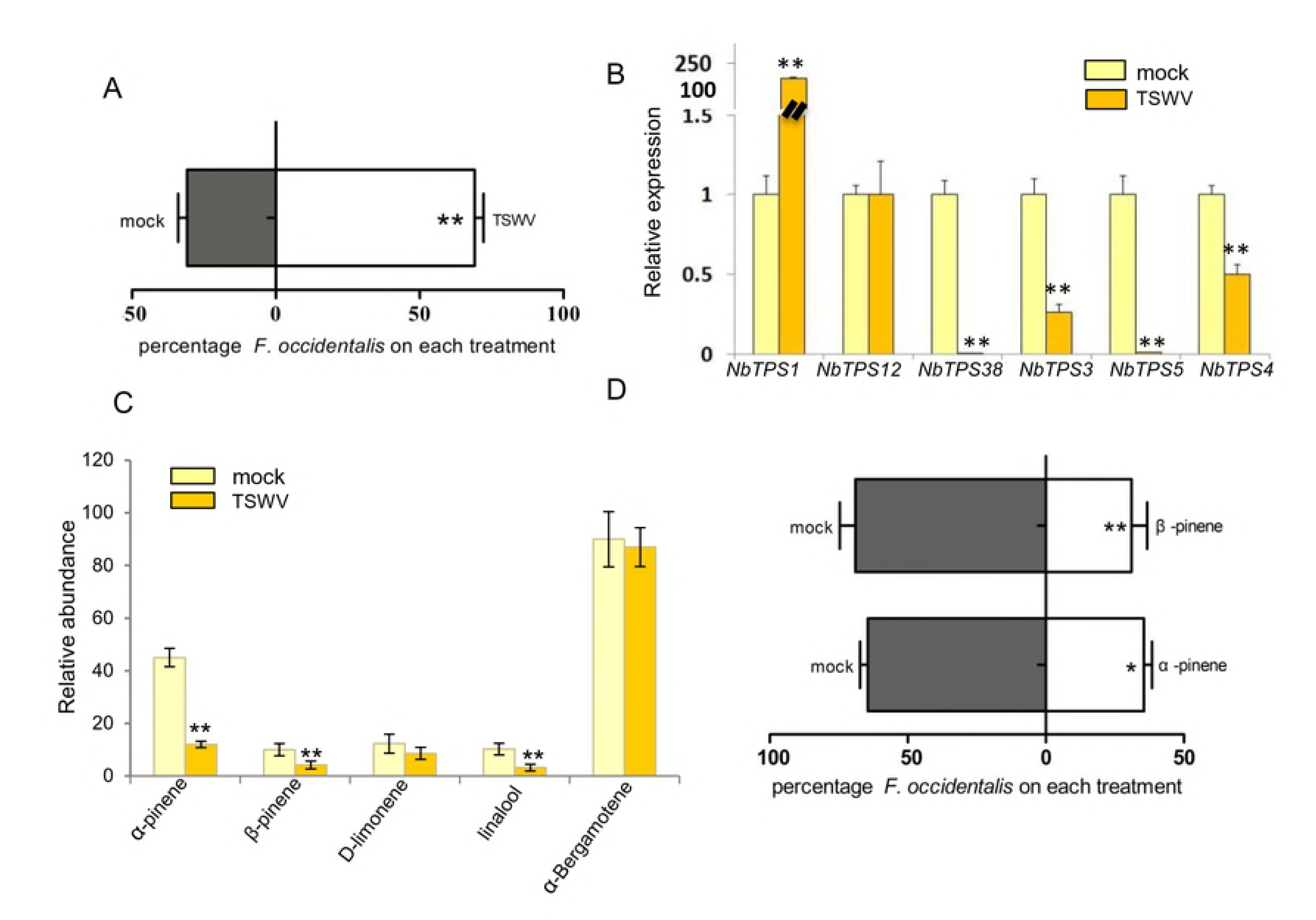
TSWV infection increases attraction to the thrips vector in a terpene-dependent manner. (A) TSWV-infected *N. benthamiana* is more attractive to Western flower thrips than the control. Three-week-old soil-cultivated *N. benthamiana* plants were infected with TSWV (TSWV) or inoculated with buffer (mock). Leaves of a similar size were used for the thrips bioassay at 14 dpi. Data are mean ± SE, n = 6. ^∗∗^P < 0.01, χ^2^-test. (B) Relative expression levels of various *TPS* genes in *N. benthamiana* after TSWV infection. Values are means ± SE, n = 5. ^∗∗^P < 0.01, Student’s *t*-test. (C) Terpenes emitted by *N. benthamiana* after TSWV infection. Values are mean relative amounts (percentage of internal standard peak area) ± SE, n = 4. ^∗∗^P < 0.01, Student’s *t*-test. (D) Monoterpene-altered TSWV-infected plants are less attractive to thrips than n-hexane control in a two-choice assay. Data are mean percentages ± SE, n = 6. ^∗∗^P < 0.01, χ^2^-test.

The attraction of insect vectors induced by the infection of other viruses is dependent on plant volatiles. We therefore measured the expression levels of *terpene synthase* (*TPS*) genes in *N. benthamiana* leaves based on our previous functional analysis of *TPS* genes [20]. Reverse-transcription quantitative PCR (RT-qPCR) analysis showed that four of the six *TPS* genes, *NbTPS3*, *NbTPS4*, *NbTPS5*, and *NbTPS38*, were significantly downregulated in TSWV-infected plants compared with the control. However, in contrast to the repressive effect of begomovirus infection on *NbTPS1* expression [20], *NbTPS1* transcript levels were ~200-fold greater in TSWV-infected plants vs. the control (Fig 1B). To explore the metabolic consequences of the altered *TPS* gene expression, we investigated changes in the emission of plant volatile compounds after TSWV infection. We applied methyl jasmonate (MeJA) to mimic WFT infestation, since this plant hormone elicits the production of various terpenes [22]. As shown in Fig 1C, five terpenes were detected in the headspace of *N. benthamiana*. Among these, the levels of three volatile monoterpenes, α-pinene, β-pinene, and linalool, significantly decreased in TSWV-infected plants compared to non-infected plants, reflecting the same trend as the downregulation of most *TPS* genes. However, there was no significant difference in the emissions of the monoterpene D-limonene or the sesquiterpene α-bergamotene (Fig 1C and S1A Fig.).

To examine whether plant monoterpenes whose levels are reduced in response to TSWV infection play a role in plant-WFT interactions, we performed two-choice assays to compare the choice of non-viruliferous WFT for these monoterpenes vs. the solvent control hexane. As expected, α-pinene and β-pinene has a repellent effect on WFT (Fig 1D).

### *NbTPS5* encodes a monoterpene synthase that produces terpenoids that are repellent to WFT

Because several TPS genes in *N. benthamiana* were severely repressed by TSWV infection, we investigated their roles in plant resistance to WFT. RT-qPCR analysis revealed that the expression levels of *TPS* genes were very low and did not notably change during the first 24 h of WFT infestation. However, the expression levels of *NbTPS5* and *NbTPS38* sharply increased at 48 h of infestation (Fig 2A). These results indicate that *NbTPS5* and *NbTPS38* function in defense responses against WFT infestation in *N. benthamiana*. We then asked whether NbTPS5 and NbTPS38 proteins are responsible for the biosynthesis of WFT repellents α-pinene and β-pinene. NbTPS38 is likely a sesquiterpene synthase, like NaTPS38 [23], as these proteins share 91% identity (S1B Fig.). Therefore, we focused on the enzyme activity of NbTPS5. We produced recombinant NbTPS5 and incubated the purified protein with its possible substrate, geranyl diphosphate (GPP), for 30 min. Gas chromatography-mass spectrometry (GC-MS) of the enzyme products showed that β-pinene and D-limonene are two monoterpene products of NbTPS5 *in vitro* (Fig 2B). These results indicate that *NbTPS5* is a key virus target gene in the host terpene biosynthesis pathway.

**Fig 2.**
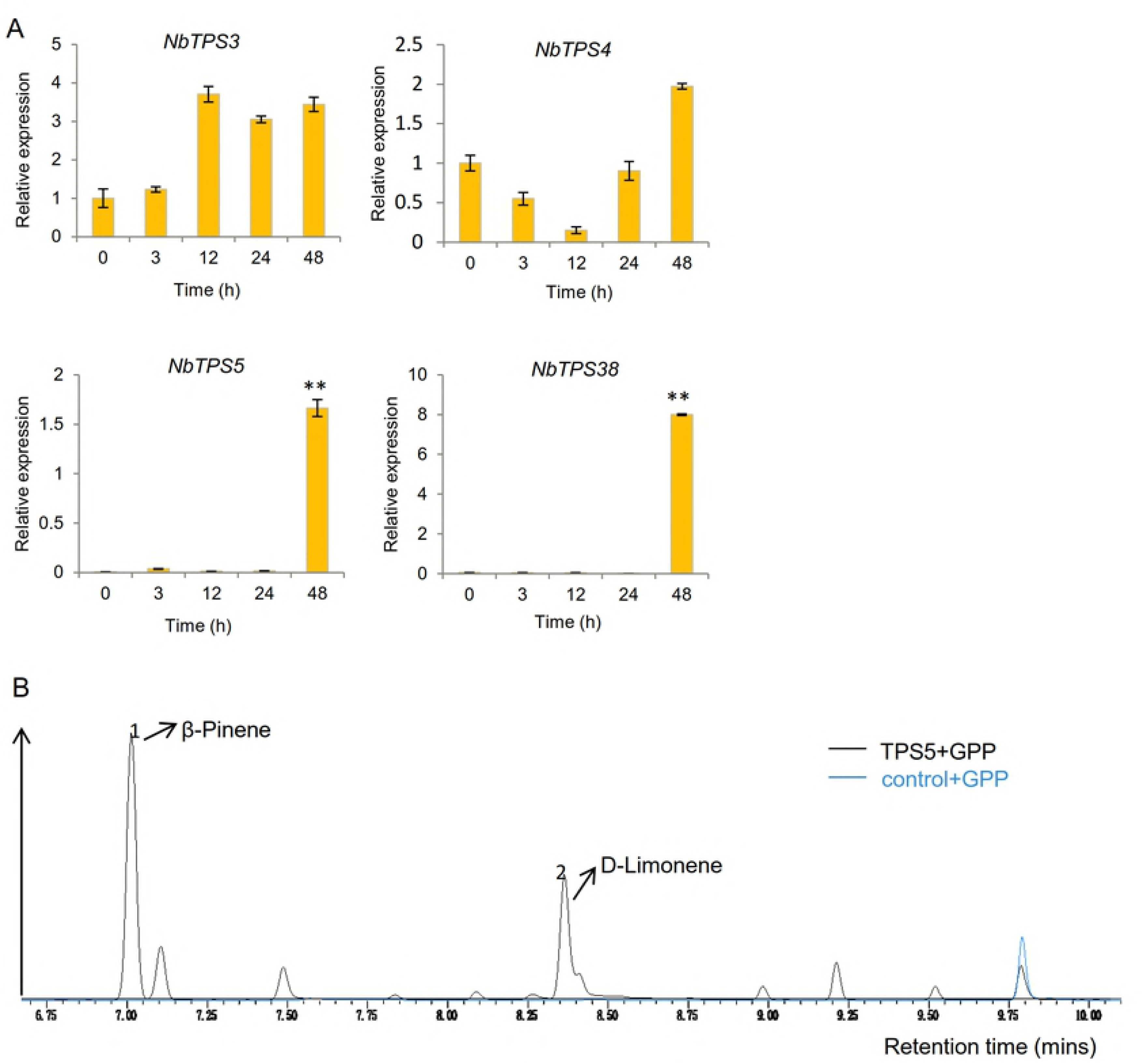
*NbTPS5* encodes a monoterpene synthase that produces repellents of WFT. (A) Relative expression level of TSWV-regulated *TPS* gene in *N. benthamiana* after thrips feeding. Four-week-old plants were grown in a single pot, and 20 adult thrips were allowed to feed on plants for 0, 3, 12, 24, or 48 h. Total RNA was prepared for RT-qPCR analysis. Values are means ± SE, n = 3. ^∗∗^P< 0.01, Student’s *t*-test. (B) NbTPS5 produces β-pinene and D-limonene *in vitro*. Gas chromatography-MS analysis of enzyme products from purified recombinant NbTPS5 and incubated with the substrate GPP. The peaks of target products are marked with arrows. The empty vector control is indicated by a blue line.

### NSs manipulates the behavior of TSWV

Our data demonstrated that the tospovirus TSWV increases the attraction of insect vector WFT to its host plant by inhibiting terpene synthase in the host. Next, to explore the protein(s) in TSWV that manipulate viral vector host choice, we selected three of the five viral proteins in TSWV, including a structural protein nucleocapsid protein (Ncp) and two non-structural proteins, NSm and NSs [12]. We used constructs harboring *CaMV35S*:*YFP* (YFP), *CaMV35S*:*YFP*-*NSs* (NSs), *CaMV35S*:*YFP*-*NSm* (NSm), and *CaMV35S*:*YFP*-*Ncp* (Ncp) in an *Agrobacterium*-mediated transient expression assay to test their possible repressive effects on *NbTPS5* expression. RT-qPCR analysis indicated that *NbTPS5* was significantly induced by NSm and Ncp but weakly repressed by NSs compared to the YFP control, suggesting that NSs mediates the repressive effect of TSWV on *NbTPS5* expression (Fig 3A). We performed a WFT two-choice assay to determine whether the expression of YFP-NSs alone is sufficient to attract WFT compared to YFP alone. As expected, without induction of other viral components (NSm or Ncp), WFT showed no significant preference for NSs-expressing leaves (Fig. 3B). We then co-expressed NSm with NSs (NSm+NSs) in *N. benthamiana* compared to NSm or NSs co-infiltrated with YFP (NSm+YFP; NSs+YFP) as the control. As shown in Fig 3C, co-expression of NSm with NSs strongly inhibited the expression of *NbTPS5* (Fig 3C). Consistent with the RT-qPCR results, plants co-expressing NSm with NSs were more attractive to WFT than control plants (Fig 3B).

**Fig 3.**
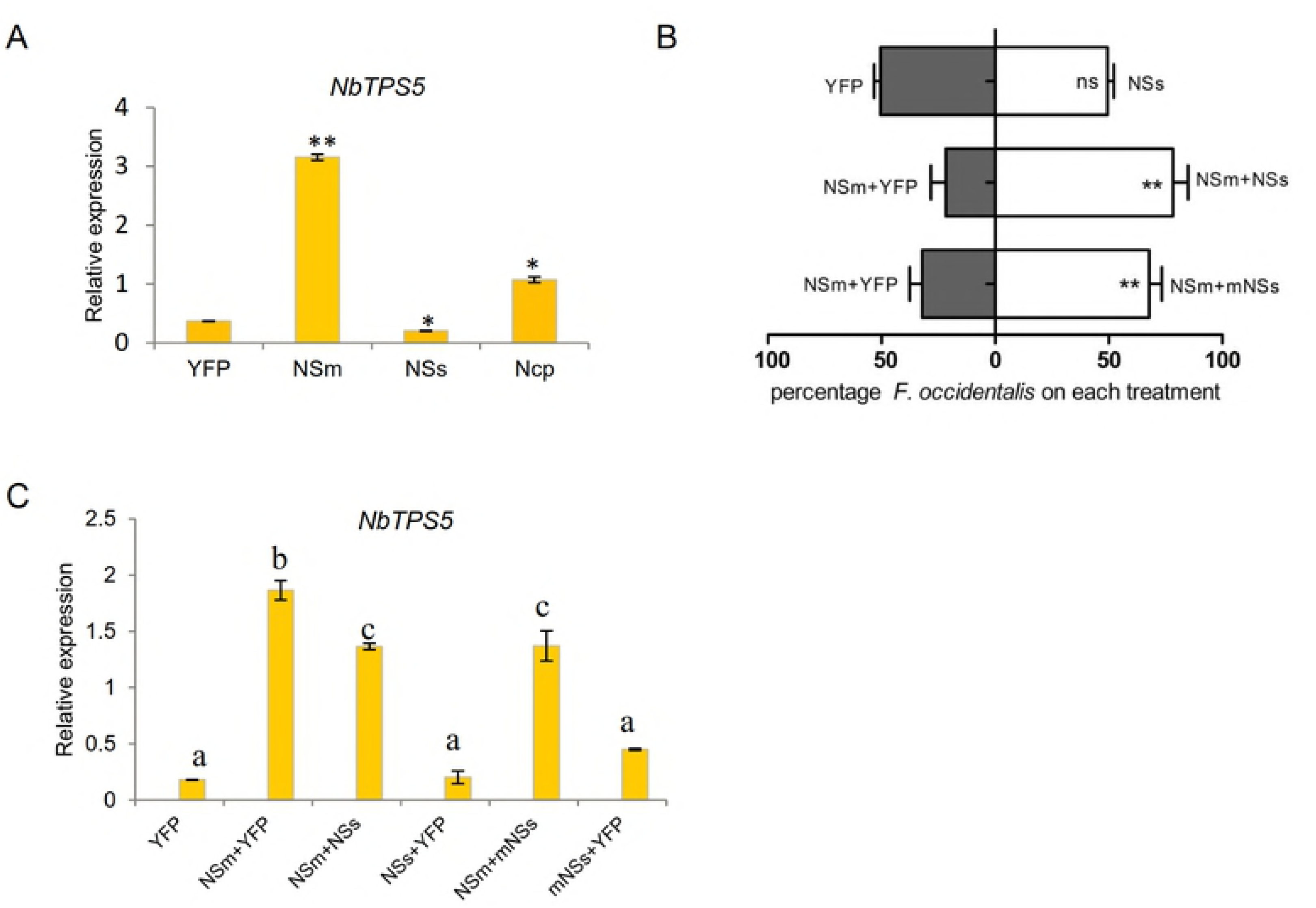
NSs from TSWV is a vector behavior manipulator. (A) Relative expression level of *NbTPS5* in *N. benthamiana* co-expressed with TSWV proteins. Four-week-old *N. benthamiana* were infiltrated with the indicated virus protein, and total RNA was prepared from the infiltrated plants at 2 dpi for RT-qPCR analysis. Values are means ± SE, n = 3. ^∗^P < 0.05, ^∗∗^P < 0.01, Student’s *t*-test. (B) Attractiveness of different infiltrated plants to thrips vector. Leaves of a similar size were used for thrips two-choice assay at 2 dpi. Data are mean percentages ± SE, n = 6. ^∗∗^P < 0.01, χ^2^-test. (C) mNSs is a mutant NSs (S48A/R51A) protein without VSR activity. Indicated virus proteins were infiltrated in *N. benthamiana* and then samples were taken for RT-qPCR analysis. Values are means ± SE, n = 3. ^∗^P < 0.05, ^∗∗^P < 0.01, Student’s *t*-test.

NSs suppresses host antiviral RNA silencing to promote TSWV infection [11]. To determine if the effect of NSs on manipulating vector behavior is dependent on its silencing suppressor activity, we explored the activity of NSs protein with an S48A/R51A mutation (mNSs) that destroys its silencing suppressor activity via RT-qPCR analysis and a WFT two-choice assay [24]. As shown in Figs 3B and 3C, mNSs retained its ability to inhibit the expression of *NbTPS5*, and plants harboring mNSs attracted more WFT than control plants (Figs 3B and 3C). Taken together, these results indicate that NSs is a viral manipulator of vector behavior that controls the attraction of WFT to the host plant in a manner independent of its virus silencing suppressor activity.

### NSs from TSWV interacts with MYC2 and its homologs MYC3 and MYC4

To explore the host protein targets of NSs, we screened an *Arabidopsis* cDNA library by yeast two-hybrid analysis and identified AtMYC2, a key components of the JA signaling pathway [25]. The JA signaling pathway plays a role in acquired resistance against thrips feeding [22]. Moreover, *NbTPS5* and *NbTPS38*, which were induced by thrips feeding, also showed a strong response to MeJA treatment (S2A Fig.). Based on the importance of the JA signaling pathway to plant-herbivore interactions, we further confirmed the interaction between AtMYC2 and NSs in a yeast cotransformation assay and a bimolecular fluorescence complementation (BiFC) assay. As shown in Fig 4A, yeast transformants harboring AD-AtMYC2 and BD-NSs could grow on SD-Leu-Trp-His medium with 0.04 mg/mL X-α-gal and turned blue, while the negative control transformants did not. These observations help confirm the AtMYC2 and NSs interaction in yeast (Fig 4A). The BiFC assay confirmed this interaction *in vivo*; as shown in Fig 4B, a strong interaction (strong fluorescence) was observed in the nucleolus, and the nucleoplasm was marked by 4’,6-diamidino-2-phenylindole (DAPI) staining (Fig 4B, first panel), while no fluorescence was observed in the negative controls (S2C Fig., first and third panels). Although AtMYC2 is localized to the nucleus [26], NSs were detected in the nucleus and cytoplasm when expressed alone (S3B Fig.). These results indicate that NSs directly interacts with and relocalizes AtMYC2 to the nucleolus, therefore disabling its activity. Since the attraction of WFT to NSs is independent of its silencing suppressor activity, we asked whether the interaction between AtMYC2 and NSs is also independent of the silencing suppressor activity of NSs. A BiFC assay of mNSs-cEYFP and AtMYC2-nEYFP provided direct evidence that mNSs interacts with AtMYC2 *in vivo* (Figs 4B, second panels), indicating that the silencing suppressor activity of NSs is independent of its interaction with AtMYC2.

**Fig 4.**
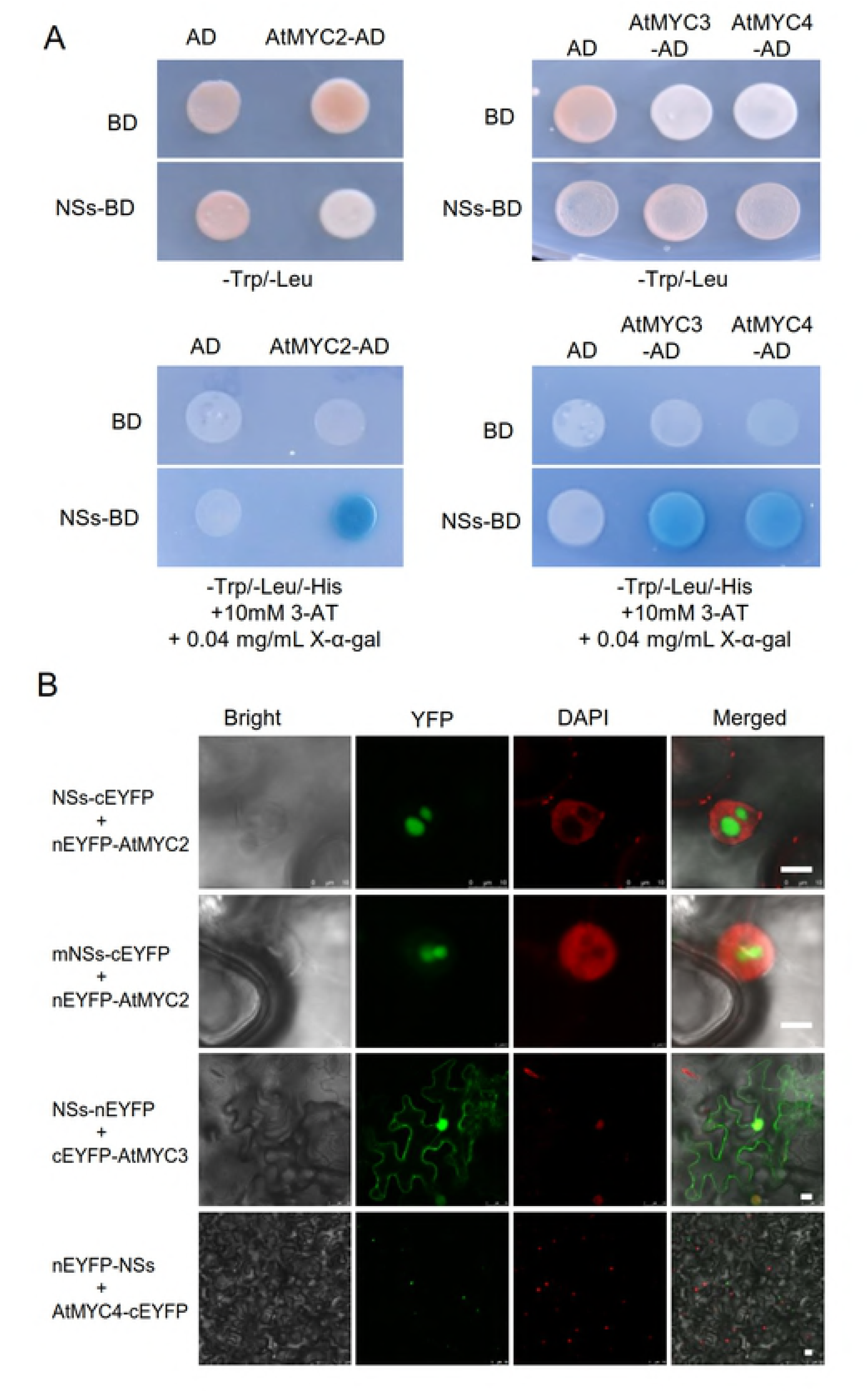
TSWV NSs interacts with MYC2 and its homologs MYC3 and MYC4. (A) Interaction between NSs and AtMYCs in a yeast two-hybrid assay. Yeast cotransformed with the indicated plasmids was spotted onto synthetic medium (SD-Leu-Trp-His) containing 0.04 mg/mL X-α-gal and 10mM 3-amino-1,2,4-triazole (3-AT). The empty vectors pGBKT7 (BD) and pGADT7 (AD) were used as negative controls. (B) Interaction between NSs/ AtMYCs and mNSs/AtMYC2 in a bimolecular fluorescence complementation assay. Nuclei in leaf epidermal cells were stained with DAPI. Bars = 10 μm.

MYC3 and MYC4 are two close homologs of bHLH transcription factors that function partially redundantly with MYC2 to activate JA responses in *Arabidopsis* [27]. To determine whether this interaction is a common feature of the MYC family during TSWV infection, we performed a yeast cotransformation assay and a BiFC assay. As expected, AD-AtMYC (MYC3 and MYC4) and BD-NSs yeast transformants turned blue when grown on SD-Leu-Trp-His medium with 0.04 mg/mL X-α-gal (Figs 4A). In the BiFC assay, *N. benthamiana* coexpressing MYC3 and NSs exhibited strong fluorescence in the membrane and nucleus (Figs 4B, third panels). Fluorescence was also detected in the cytoplasm in plants coexpressing MYC4 and NSs (Figs 4B, fourth panels). We further examined the interaction with the bHLH transcription factor NbMYC2, a homologous protein of AtMYC2 in *N. benthamiana* by yeast cotransformation and BiFC assay [20,28]. NbMYC2 could grow on the selection medium and interacted with NSs in an area of the cytoplasm near the nucleus (S3A Fig.). TSWV represents the American- and *Tomato zonate spot virus* (TZSV) represents the Euro/Asian-type tospoviruses. We further confirmed this conserved interaction between TZSV NSs and AtMYC2 (S4 Fig.) [29].

The similarity of these protein interactions prompted us to identify the NSs domain that interacts with plant MYCs. We constructed three truncated mutant AtMYC2 proteins and examined their interactions with TSWV NSs in a BiFC assay. NSs interacted with the middle (MID) domain or bHLH domain of AtMYC2 (S4A Fig.); similar results were obtained with mNSs (S4B Fig.). Taken together, these results demonstrate that NSs targets MYC2 through the bHLH or MID domain, which is not involved with its silencing suppressor activity.

### MYCs regulates volatile-dependent immunity against WFT in *Arabidopsis*

MYC2 plays an important role in JA-regulated plant defense responses, including plant volatile biosynthesis, and it directly regulates *TPS10* transcript levels [20,30,31]. Thus, we hypothesized that AtMYC2, which interacts with virulence factor NSs, is involved in the viral-induced, volatile-dependent attraction of WFT to the host plant. To validate this hypothesis, we performed a GUS staining assay using two transgenic *Arabidopsis* lines expressing an *AtMYC2* or *AtTPS10* promoter:*GUS* reporter gene. As shown in Fig 4A, high *GUS* expression was detected after 24 h of WFT feeding. This expression pattern suggests that *AtMYC2* and *AtTPS10* both function in defense responses against WFT in *Arabidopsis* (Fig 5A).

**Fig 5.**
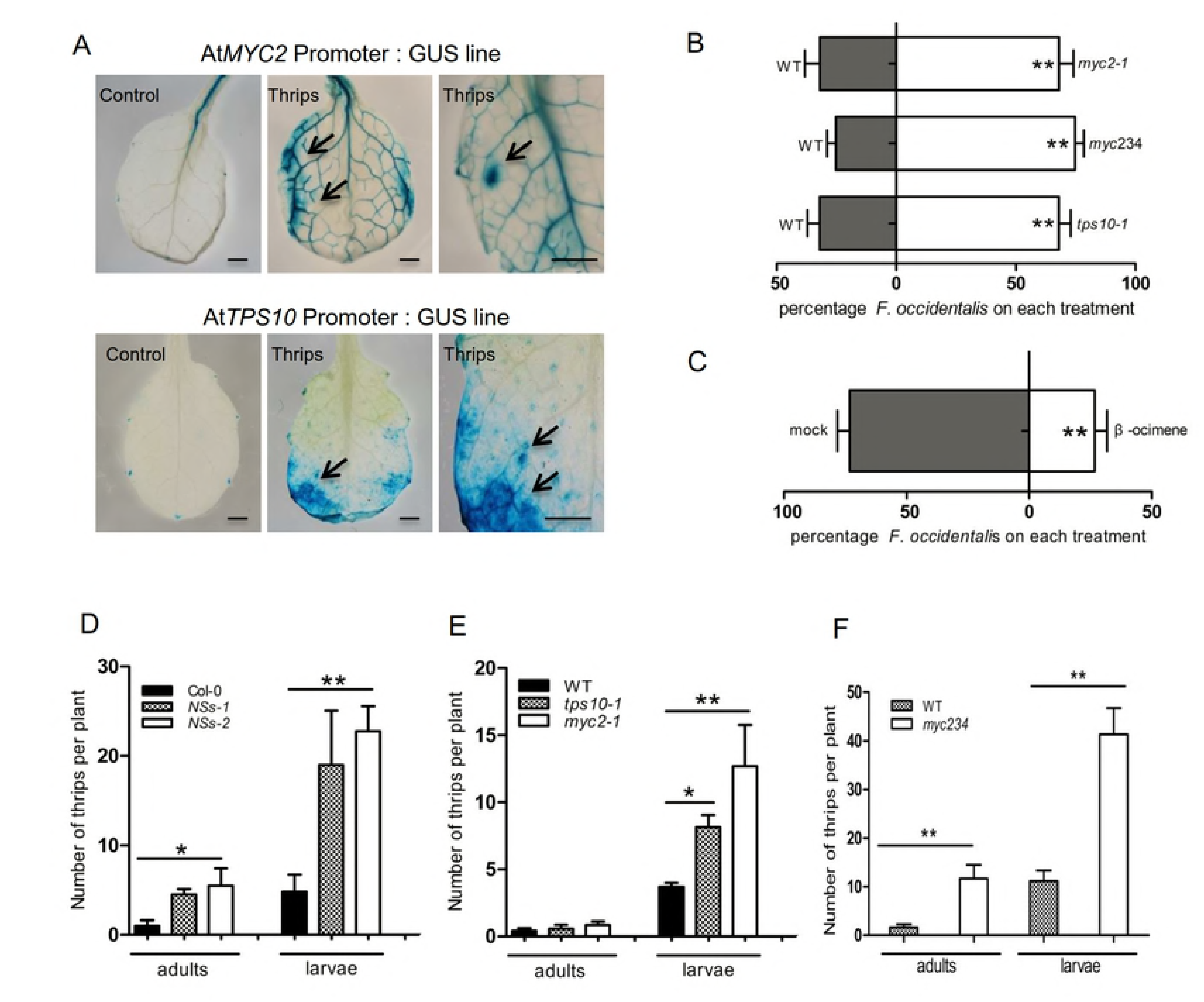
MYC2 and its homologs are essential regulators of host immunity responses against WFT. (A) GUS staining of *AtMYC2p*-*GUS* and *AtTPS10p*-*GUS* seedlings after 24 h of thrips feeding. An untreated line was used as a control. Arrows indicate thrips feeding sites. Bars = 2 mm. (B) *myc2*-*1*, *myc234*, and *tps10*-*1* mutants are more attractive to thrips than wild type. Three-week-old *Arabidopsis* plants cultured in MS medium were used for the thrips two-choice assay. Data are mean percentages ± SE, n = 6. ^∗∗^P < 0.01, χ^2^-test. (C) β-ocimene is less attractive to thrips than mock treatment in a two-choice assay. Data are mean percentages ± SE, n = 6. ^∗∗^P < 0.01, χ^2^-test. (D-F) The effects of different genes on the number of thrips. Seven adult females fed on each three-week-old *Arabidopsis* line. After 2 weeks, new larvae and adults were counted. Values are means ± SE, n = 8. ^∗^P < 0.05, ^∗∗^P < 0.01, Student’s *t*-test.

To analyze the effects of *AtMYC2* and *AtTPS10* on the feeding preferences of thrips, we performed two-choice assays using *myc2*-*1*, *tps10*-*1*, and wild-type Col-0 *Arabidopsis.* As shown in Fig 5B, the *myc2*-*1* and *tps10*-*1* mutants were more attractive to WFT than wild type. We also tested the effect of *myc234* on host preference, finding that WFT preferred to feed on this mutant over wild type (Fig 5B). *AtTPS10* encodes a monoterpene synthase, which produces β-ocimene and small amounts of cyclic monoterpenes [32]. We therefore carried out a two-choice assay of β-ocimene to examine whether the attraction of *tps10* is terpene-dependent. As expected, β-ocimene had a strong repellent effect on WFT (Fig 5C). These results indicate that *AtMYC2* is essential for terpene-dependent immunity against the thrips vector.

TSWV increases the population density of the next generation of WFT [22]. We investigated whether this effect is due to the deactivation of AtMYCs by NSs by performing a modified thrips spawning experiment [21]. Seven female adults were allowed to feed on *35S:YFP*-*TSWV NSs* (*NSs*-*1*; *NSs*-*2*) or wild-type *Arabidopsis* for two weeks. We counted the number of new adults and larvae to analyze the effect of NSs on the thrips population. Plants expressing *NSs* were more suitable for WFT population growth than wild type (Fig 5D). We reasoned that NSs targets MYCs to disable the activation of *terpene synthase* genes, thereby attenuating the defense of the host plant against thrips. To investigate this hypothesis, we conducted another spawning experiment using *myc2*-*1*, *tps10*-*1*, and *myc234* mutants. More WFT were found on the mutants compared with wild type; these lines were equally suitable for WFT growth compared to the lines expressing *NSs*, confirming the important role for NSs in the tripartite WFT-TSWV-plant interaction (Figs 5E and 5F). In summary, our results suggest that NSs targets MYCs to attenuate host defense responses to thrips, thereby manipulating terpene-dependent chemical communication between the plant and thrips vector.

## Discussion

### TSWV suppresses host terpene biosynthesis and promotes the performance of its thrips vector

Two types of TSWV-infected plants, *Arabidopsis* and pepper, have been shown to attract more WFT vectors than non-infected plants [7,21]. In this study, we demonstrated the existence of virus-induced manipulation of *N. benthamiana*, another TSWV host (Fig 1A). We demonstrated that TSWV manipulates this process to affect the behavior of the WFT vector. The volatile terpenoids emitted by virus-infected *N. benthamiana* plants contained fewer repellents to WFT than healthy plants (Fig 1C). *Arabidopsis* displayed similar levels of terpene-dependent attraction to WFT (Fig 5C). These consistent results in different species indicate that this feature is common among various TSWV hosts.

Instead of increasing the level of attractant, TSWV represses the production of various monoterpene repellents for WFT, e.g., α-pinene and β-pinene in *N. benthamiana* and β-ocimene in *Arabidopsis*. The decline in repellent levels has strong benefits for WFT, despite the possible negative physiological consequences of carrying TSWV, and therefore, WFT prefers to attack virus-infected plants. Linalool, another monoterpene whose production was also repressed by TSWV infection in our study (Fig. 1C), inhibits the growth of WFT [33,34]. We found that the production of the monoterpene synthase, NbTPS5, was greatly suppressed by TSWV infection. This enzyme synthesizes monoterpene repellents, thereby attracting the WFT vector. Intriguingly, the basal expression level of *NbTPS5* was extremely low, and it was highly induced in response to other essential TSWV components, i.e., NSm and Ncp, pointing to an arms race between defense and counter-defense responses in the TSWV-host interaction (Fig 1D and Fig 2B). Moreover, TSWV infection reduced the expression of various *TPSs*, including *NbTPS5*. In addition to *NbTPS5*, *NbTPS38* was also downregulated by TSWV infection (Fig 1B). Interestingly, the likely sesquiterpene synthase gene *NbTPS38* was downregulated in response to NSm and Ncp, as well as NSs, supporting the notion that multiple proteins contribute to the complex tripartite TSWV-WFT-plant interaction (S2B Fig.). The expression of *MYC2*-*TPS10*, which functions in the monoterpene defense pathway in *Arabidopsis*, was also attenuated in response to TSWV to benefit WFT (Fig 5) [20,32]. The conserved strategy of suppressed monoterpene biosynthesis is utilized by TSWV in different plants, suggesting that terpenoid biosynthesis is an evolutionarily conserved target for plant viruses to promote vector performance.

### MYCs are involved in virus-induced terpene-dependent attraction to the insect vector

In addition to the terpene biosynthesis pathway, anti-herbivore phytohormone pathways can be regulated to increase the fitness of the host plant to vectors [35–40]. Numerous studies have described the importance of JA in plant defense responses against pathogen and insect attack [22,41,42]. JA-dependent plant defense control WFT performance and preference, and TSWV infection reduces the levels of these responses. In JA-insensitive *coi1*-*1* mutants, WFT did not show a preference for TSWV-infected plants [9]. The current results suggest that the MYC proteins involved in the JA pathway are responsible for plant terpene immunity for WFT (Fig 5A-C). *MYCs* are downstream genes of *COI1*, and MYC2-orchestrated transcriptional reprogramming is a central theme of JA signaling [25]. Functional blocking of *MYCs* increases WFT preference and promotes WFT performance, including developmental duration and fecundity (Fig 5D-F). The antagonistic relationship between JA and salicylic acid signaling facilitates interactions between the thrips vector and TSWV [21]. Our results indicate that salicylic acid also mediates *NPR1*-indepedent immune responses to thrips (S5 Fig.). We hypothesize that several MYC-regulated indole and aliphatic glucosinolates that function as defensive chemicals against herbivores might be repressed, as is the case for βC1 in begomovirus [43]. Alternatively, the levels of nutrients (such as amino acids) are likely altered in the host, which could affect the feeding behavior and preference of thrips, as previously reported [44].

There is some evidence that plant viruses target JA signaling in several phytopathological systems. For example, we previously demonstrated that βC1 of *Tomato yellow leaf curl China virus* (TYLCCNV) interacts with MYC2 to subvert plant resistance and to promote vector performance [20]. Similarly, 2b of *Cucumber mosaic virus* also suppresses JA signaling, and *myc234* triple mutant plants were also observed to induce the attraction of the CMV aphid vector [20,45]. Importantly, the same strategy is employed by different viruses, suggesting that it is a general feature of tritrophic virus-vector-plant interactions. However, each virus exhibits specific features throughout its lifecycle and therefore, the details of the interactions vary. For instance, βC1 of TYLCCNV alone is sufficient to attract the whitefly vector, but NSs of TSWV is not, as it requires the involvement of another elicitor such as NSm (Fig 3B).

### A novel function of NSs related to vector attraction

TSWV NSs is an avirulence determinant that triggers a hypersensitive response in resistant plants [46]. NSs is also a well-known viral suppressor of host RNA interference in both plants and insects and is essential for TSWV transmission by WFT [5,11,13]. Here, we identified NSs as a viral effector that attenuates the host defense system by suppressing MYC proteins (Fig 4B). The expression of *NSs* in *Arabidopsis* is sufficient to promote WFT performance (Fig 5D). Interestingly, the vector manipulation behaviors of both 2b of *Cucumber mosaic virus* and NSs of TSWV are independent of their silencing suppressor activity (Fig 3B and Fig 4B) [45], these two cellular activities of viral proteins may function in parallel in plant cells. NSs, a multifunctional protein, employs other strategies to control vector attraction not related to its silencing suppressor activity, e.g., the relocation of MYC proteins to downregulate JA signaling.

### Bunyaviruses play conserved roles in manipulating the behaviors of insect vectors

By interrupting MYC-regulated plant defense via NSs, TSWV indirectly manipulates the preference and performance of WFT. This type of behavioral manipulation has also been observed in animal-infecting bunyaviruses. As early as 1980, *La Crosse virus* (LACV) was reported to modify the feeding behavior of mosquito vectors [2]. *Rift Valley fever virus* (RVFV) was found to affect mosquito vector morbidity and mortality [3]. These studies revealed a conserved trait within members of *Bunyavirales*. However, the molecular mechanism underlying this manipulation was unclear, and no specific information was available regarding viral determinants of the virus-host-vector interaction in other bunyaviruses. Our study identified NSs of TSWV as the first vector behavior manipulator that suppresses host plant defense responses to attract and benefit the fitness of WFT, which in turn facilitates disease dispersal plant to plant. This provides a potential strategy for bunyaviruses-induced changes to vectors, extending our understanding of this tritrophic interaction at the molecular level. However, many questions remain to be answered before we can fully understand this complex tritrophic interaction.

## Materials and Methods

### Plant materials

*Nicotiana benthamiana* and *Arabidopsis thaliana* (Col-0) plants were grown in insect-free growth chambers following standard procedures [20]. The *Arabidopsis myc2*-*1*, *tps10*-*1*, and *myc234* mutants (Col-0 background) were described previously [20]. The *35S:YFP*-*NSs* transgenic lines *NSs*-*1* and *NSs*-*2* were generated using the *Agrobacterium*-mediated floral-dip method [41].

### Naive Western flower thrips colony and mechanical inoculation of TSWV

A starting colony of Western flower thrips (WFT, *Frankliniella occidentalis*) (Pergande) (*Thysanoptera*: *Thripidae*) was kindly provided by Prof. Youjun Zhang (Institute of Vegetables and Flowers, Chinese Academy of Agricultural Sciences). The thrips were maintained on *Phaseolus vulgaris* in a climate chamber as described previously [47]. *Tomato spotted wilt orthotospovirus* (isolate TSWV-YN) was isolated from Yunnan, China, and amplified by mechanically inoculation onto *N. benthamiana* as described by Mandal et al. [48]. Infected leaves were ground in 0.05 M phosphate buffer (pH 7.0) and applied to the host plant using a soft finger-rubbing technique. Infected plants were tested at 10-14 dpi by RT-qPCR prior to the thrips two-choice assays.

### Thrips two-choice assay

The two-choice assay was performed as described previously [44]. Petri dishes 16 cm in diameter were prepared by covering with moist filter paper. For *N. benthamiana*, detached leaves of TSWV-infected plants (TSWV) and non-infected plants (mock) were separately placed in a Petri dish. Fifty *F. occidentalis* adults were released to the center of the Petri dish containing two leaves. The number of thrips that settled on each leaf was counted at 24 h after release. For *Arabidopsis*, plants were cultivated on solid Murashige and Skoog medium for 3 weeks, and whole plants were used for the two-choice assay. For two-choice assays using monoterpene, 2 cm × 2 cm filter paper containing 40 μL of a 1:100 (v/v) solution of standard chemical substance from Sigma in n-hexane or n-hexane alone (as a control) were placed in a Petri dish. Thrips were released between the two tested samples, and the thrips were counted 5 min after release. The Petri dishes were contained in a thrips culture chamber throughout the experiment to maintain consistent growth conditions.

### Thrips infestation assay

Plants were infested with non-viruliferous thrips as described previously [22]. Twenty adults (7-14 d after eclosure) were grouped and starved for 3 h before the plant infestation assay. *Arabidopsis* plants grown on solid MS medium or soil-grown *N. benthamiana* plants were infested with adult thrips for the indicated time period. The thrips were gently removed and the leaf samples collected in liquid nitrogen for further analysis. For the *GUS*-reporter line expression assays, transgenic *Arabidopsis* plants were infested with thrips for 24 h, followed by GUS activity analysis. The experiment was repeated at least twice with similar results.

### Volatile analysis

The collection, isolation, and identification of volatiles from *N. benthamiana* plants were performed as described previously [20,49]. Volatiles emitted from TSWV-infected and healthy plants were collected for 12 h at a gas flow rate of 300 mL/min and analyzed by gas chromatography. At least four plants per group were used.

### Terpene synthase enzyme activity

The complete open reading frame of *NbTPS5* was cloned into the *pGST*-*DC* vector. The protein was expressed in *E. coli* strain BL21, and purified GST protein was used as a negative control. Protein purification was conducted as previously described [23]. Recombinant NbTPS5 or control GST protein was incubated with the substrate, geranyl diphosphate (GPP), in reaction buffer (pH 7.5, 5 mM Tris-HCl, 5% v/v glycerol, 10 mM MgCl_2_, 2 mM dithiothreitol) at 30°C for 30 min. Enzyme activity was measured using a solid phase microextraction (SPME; Supelco, Belafonte, PA, USA) fiber consisting of 100 μm polydimethylsiloxane (Supelco). Chemical analysis was performed by gas chromatography-mass spectrometry (GC-MS) (Shimadzu, QP2010). GC was performed with a DB5MS column (Agilent, Santa Clara, CA, USA, 30 m × 0.25 mm × 0.25 μm). The SPME fiber was thermally desorbed in the injector at 250°C for 1 min. The initial oven temperature was held for 3 min, increased to 240°C with a gradient of 5°C/min, and maintained at 240°C for 5 min. The inlet temperature was 270°C.

### *Agrobacterium*-mediated transient expression

Constructs harboring *YFP*-*NSs*, *YFP*-*NSm*, and *YFP*-*Ncp* were generated by PCR cloning of the coding sequences of *NSs*, *NSm*, and *Ncp* of TSWV-YN into vector *pH7*-*YFP*-*DC*, respectively, using Gateway technology (Life Technologies). PCR-based mutagenesis was used to construct the NSs (S48A/R51A) mutant with the primers listed in S1 Table. *Agrobacterium*-mediated transient expression assays were performed by agroinfiltration of *Agrobacterium* carrying the binary plasmids into the the basal (abaxial) sides of *N. benthamiana* leaves [24]. Infiltrated leaves were collected at 2 dpi and frozen in liquid nitrogen for RT-qPCR. Plants infiltrated with empty vector (*pH7.WG2Y*) were used as a negative control. The experiment was repeated at least twice with similar results.

### Yeast two-hybrid analysis

Full-length *NSs* was amplified and inserted into the pGBT9 vector. The *Arabidopsis* Mate and Plate Library was screened as previously described [20]. The interaction between NSs and MYCs was confirmed according to the manufacturer’s protocol (Clontech). The pGBT9-NSs and pGAD424-MYC constructs were co-transformed into yeast strain Y2HGold. Yeast cotransformed with the indicated plasmids was spotted onto synthetic medium (SD-Leu-Trp-His) containing 10mM 3-amino-1,2,4-triazole and 0.04 mg/mL X-α-gal. The empty vectors pGBKT7 (BD) and pGADT7 (AD) were used as negative controls.

### Bimolecular fluorescence complementation (BiFC)

BiFC was performed as described previously [20,50]. The indicated constructs were infiltrated in four-week-old *N. benthamiana* leaves via agroinfiltration. The fluorescent signals and DAPI staining were detected at 2 dpi via confocal microscopy.

### Quantitative RT-PCR

Total RNA was extracted from leaf and plant samples using an RNeasy Plant Mini Kit (Qiagen) with column DNase treatment [51]. RNA was reverse transcribed using TransScript One-Step gDNA Removal and cDNA Synthesis SuperMix (TransGen Biotech, China). Four to six independent biological samples were collected and analyzed. RT-qPCR was performed using SYBR Green Real-Time PCR Master Mix (Toyobo, China) on the CFX 96 system (Bio-Rad). *Arabidopsis Actin*-*2* and *N. benthamiana EF1α* were used as the internal controls [22].

### Thrips spawning assay

The thrips spawning assay was performed as described previously with some modifications [21]. *Arabidopsis* plants were grown in soil covered with Parafilm (Bemis, USA) to prevent any thrips from escaping and to facilitate counting. Three-week-old plants were placed in an acryl cylinder chamber (7 cm × 5 cm) and covered with a fine mesh. Seven female adults (7-14 d after eclosure) were allowed to infest a single plant for two weeks, and new larvae and adult thrips were counted. Eight plants of each genotype were used per experiment. The experiment was repeated at least twice with similar results.

### GUS staining

Transgenic *Arabidopsis* plants expressing *AtMYC2* or the *AtTPS10* promoter:*GUS* reporter gene were infested with thrips for 24 h and incubated in GUS staining buffer (0.5 mg/mL X-glucuronide, 0.5 mM potassium ferricyanide, 0.5 mM potassium ferrocyanide, 10 mM EDTA, 0.1% Triton X-100, 0.1 M pH 7.0 phosphate buffer) at 37°C overnight. The stained seedlings were cleared by washing with 70% ethanol. Untreated plants were used as a negative control. The experiment was repeated at least twice with similar results.

### Data analysis

Significant differences in gene expression and volatile organic compound levels were determined by Student’s t tests or one-way ANOVA; if the ANOVA result was significant (P < 0.05), Duncan’s multiple range tests were used to detect significant differences between groups. Thrips choices between different treatments were analyzed by Pearson χ^2^ tests. All statistical tests were carried out with GraphPad Prism.

### Accession numbers

Sequence data in this study can be found in GenBank/EMBL or TAIR (www.Arabidopsis.org) under the following accession numbers: AtMYC2 (At1g32640), AtMYC3 (AT5G46760), AtMYC4 (AT4G17880), AtTPS10 (At2g24210), NbTPS1 (KF990999), TSWV NSs (JF960235.1), TSWV NSm (JF960236.1), and TSWV Ncp (JF960235.1).

## Acknowledgments

We thank Profs. Youjun Zhang (Chinese Academy of Agricultural Sciences, China), and Shujun Wei (Beijing Forest and Agriculture Academy of Sciences, China) for providing the WFT clones. We also thank Prof. Guodong Wang for providing GPP (Institute of Genetics and Developmental Biology, Chinese Academy of Sciences). This study was supported by the Chinese Academy of Sciences (Strategic Priority Research Program Grant NO. XDB11040300) and National Natural Science Foundation of China (31522046, 31672001, 31830073).

## Supporting information

**S1 Fig. Effect of TSWV infection on terpenoid biosynthesis in *N. benthamiana*.**

(A) Terpenes emitted by *N. benthamiana* after TSWV infection. (B) Sequence alignment of NbTPS38 and NaTPS38 (a sesquiterpene synthase in *Nicotiana attenuata*). NbTPS38 and NaTPS38 share 91% protein sequence similarity.

**S2 Fig. JA-responsive *NbTPS5* and *NbTPS38* expression.**

(A) Relative *NbTPS5* and *NbTPS38* expression levels in *N. benthamiana* after MeJA treatment. Total RNA was prepared from plants at 24 h after MeJA treatment for RT-qPCR analysis. Values are means ± SE, n = 3. ^∗∗^P< 0.01, Student’s *t*-test. (B) Relative expression of *NbTPS38* in *N. benthamiana* after challenge with viral protein. Values are means ± SE, n = 3. ^∗^P < 0.05, ^∗∗^P < 0.01, Student’s *t*-test. (C) Negative control in the bimolecular fluorescence complementation assay. Nuclei in leaf epidermal cells were stained with DAPI. Bars = 25 μm.

**S3 Fig. NSs interacts with and relocalizes MYC2 and its homologs.**

(A) Interaction between TSWV NSs and NbMYC2 in yeast cotransformation and bimolecular fluorescence complementation assay. Nuclei in leaf epidermal cells were stained with DAPI. Bars = 10 μm. (B) Subcellular localization of AtMYC2 and TSWV NSs. YFP-AtMYC2 and YFP-NSs were expressed in *N. benthamiana* via *Agrobacterium*-mediated transient infiltration. Typical cells were observed by confocal microscopy.

**S4 Fig. NSs interacts with the middle and bHLH domains of MYC2.**

(A-B) Bimolecular fluorescence complementation assay to identify the interaction domain of AtMYC2 with NSs and mNSs. Three deletion derivatives of AtMYC2 are shown in the schematic diagram on the right. Nuclei in the epidermal cells on the left were stained with DAPI. Bars = 25 μm. MID, middle domain.

**S5 Fig. Salicylic acid-mediated *NPR1*-indepedent defense response to thrips.**

Seven adult females were infested on salicylic acid (*nahg* or *npr1*) mutants for 2 weeks, and new larvae were counted. Values are means ± SE, n = 8. ^∗^P < 0.05, ^∗∗^P < 0.01, Student’s *t*-test.

**S6 Fig. NSs from Tospoviruses interacts with AtMYC2.**

(A) Alignment of NSs from American- and Asian-type tospoviruses by ClustalW. TSWV, *Groundnut ring spot virus* (GRSV) and *Impatiens necrotic spot virus* (INSV) are from American-type tospoviruses while TZSV and *Watermelon silver mottle virus* (WSMoV) are from Euro/Asian-type tospoviruses. (B) Interaction between TZSV NSs and AtMYC2 in bimolecular fluorescence complementation assay. Nuclei in leaf epidermal cells were stained with DAPI. Bars = 10 μm.

**S1 Table.**
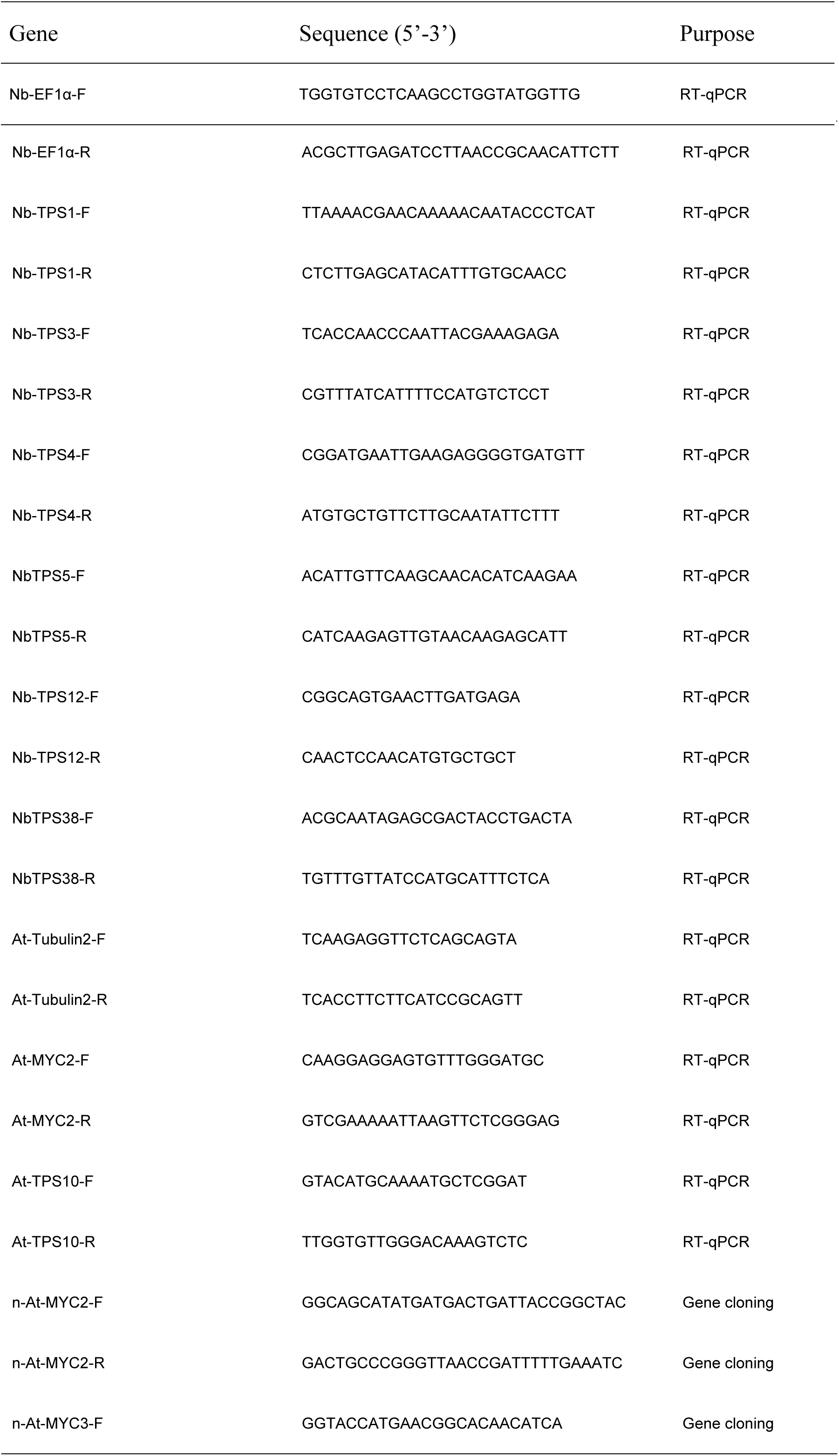
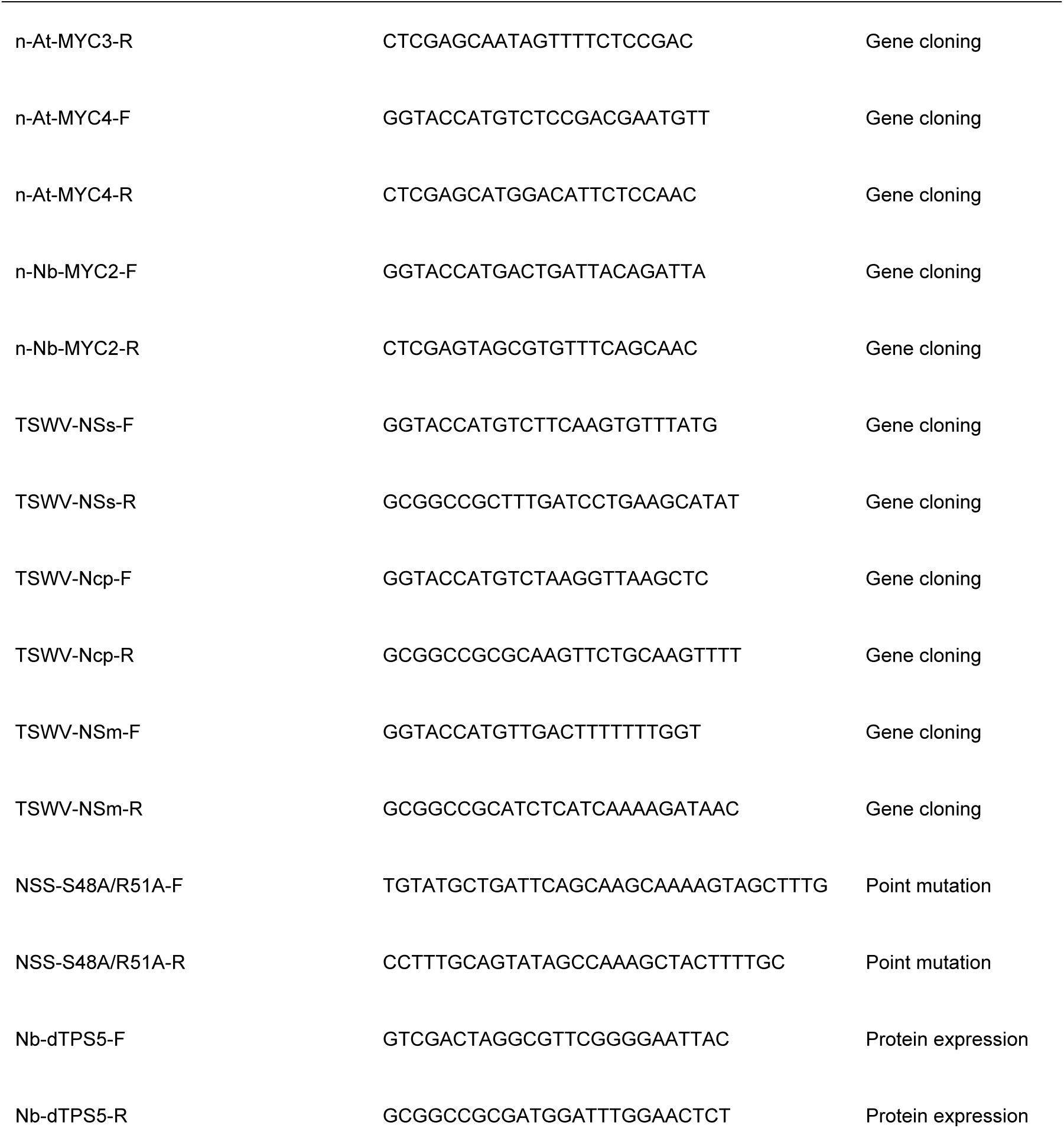
DNA primers used in this study.

